# Role of oxidation of excitation-contraction coupling machinery in age-dependent loss of muscle function in *C. elegans*

**DOI:** 10.1101/2021.12.05.471286

**Authors:** Haikel Dridi, Frances Forrester, Alisa Umanskaya, Wenjun Xie, Steven Reiken, Alain Lacampagne, Andrew R. Marks

**Affiliations:** Department of Physiology and Cellular Biophysics, The Clyde and Helen Wu Center for Molecular Cardiology, Department of Medicine, Columbia University Vagelos College of Physicians and Surgeons New York, NY, USA; PhyMedExp, Montpellier University, INSERM, CNRS, CHRU Montpellier, 34295, Montpellier, France; Medical Intensive Care Unit, Montpellier University and Montpellier University Health Care Center, 34295, Montpellier, France

## Abstract

Age-dependent loss of body wall muscle function and impaired locomotion occur within 2 weeks in *C. elegans*; however, the underlying mechanism has not been fully elucidated. In humans, age-dependent loss of muscle function occurs at about 80 years of age and has been linked to dysfunction of ryanodine receptor (RyR)/intracellular calcium (Ca^2+^) release channels on the sarcoplasmic reticulum (SR). Mammalian skeletal muscle RyR1 channels undergo age-related remodeling due to oxidative overload, leading to loss of the stabilizing subunit calstabin1 (FKBP12) from the channel macromolecular complex. This destabilizes the closed state of the channel resulting in intracellular Ca^2+^ leak, reduced muscle function, and impaired exercise capacity. We now show that the *C. elegans* RyR homolog, *UNC-68*, exhibits a remarkable degree of evolutionary conservation with mammalian RyR channels and similar age-dependent dysfunction. Like RyR1 in mammals *UNC-*68 encodes a protein that comprises a macromolecular complex which includes the calstabin1 homolog FKB-2 and is immunoreactive with antibodies raised against the RyR1 complex. Further, as in aged mammals, *UNC-68* is oxidized and depleted of FKB-2 in an age-dependent manner, resulting in “leaky” channels, depleted SR Ca^2+^ stores, reduced body wall muscle Ca^2+^ transients, and age-dependent muscle weakness. FKB-2 *(ok3007)-*deficient worms exhibit reduced exercise capacity. Pharmacologically induced oxidization of *UNC-68* and depletion of FKB-2 from the channel independently caused reduced body wall muscle Ca^2+^ transients. Preventing FKB-2 depletion from the *UNC-68* macromolecular complex using the Rycal drug S107 improved muscle Ca^2+^ transients and function. Taken together, these data suggest that *UNC-68* oxidation plays a role in age-dependent loss of muscle function. Remarkably, this age-dependent loss of muscle function induced by oxidative overload, which takes ~2 years in mice and ~80 years in humans, occurs in less than 2-3 weeks in *C. elegans*, suggesting that reduced anti-oxidant capacity may contribute to the differences in life span amongst species.

## INTRODUCTION

Approximately 50% of humans over the age of 80 develop muscle weakness, which contributes to falls and hip fractures, a leading cause of mortality in the elderly (1–3). Strikingly, despite an approximately 2,000-fold difference in the lifespans of humans and *C. elegans* (4, 5), both exhibit oxidative overload induced age-dependent reductions in muscle function and motor activity that ultimately contribute to senescence and death. Due to its short lifespan and well-characterized genome, *C. elegans* has been used as a model to study the genetics of aging and lifespan determination (6, 7), including the age-dependent decline in locomotion (4, 8). Age-dependent reduction in locomotion in *C. elegans* has been attributed to degeneration of the nervous system (9) and the body wall musculature (10). Here we investigated the role of the ryanodine receptor/Ca^2+^ release channel (RyR) homolog, *UNC-68,* in age-dependent loss of muscle function in *C. elegans*.

Mammalian RyR1 is the major intracellular Ca^2+^ release channel in skeletal muscle required for excitation-contraction (E-C) coupling (11, 12). Peak intracellular Ca^2+^ transients evoked by sarcolemmal depolarization decrease with age (13), and this decrease is associated with a reduced SR Ca*^2+^* release (14) that directly determines the force production of skeletal muscle. Our group has shown that a mechanism underlying age-dependent loss of muscle function is RyR1 channel oxidation which depletes the channel complex of the stabilizing subunit calstabin1 (calcium channel stabilizing binding protein type 1, or FKBP12), resulting in intracellular Ca^2+^ leak and muscle weakness (15, 16). RyR1 is a macromolecular complex comprised of homotetramers of four ~565 kDa RyR monomers (12, 17). Cyclic AMP (cAMP) dependent protein kinase A (PKA) (18), protein phosphatase 1 (19), phosphodiesterase PDE4D3 (20), the Ca^2+^-dependent calmodulin kinase II (CaMKII) (21, 22), and calstabin1 (23), are components of the RyR1 macromolecular complex (24). Calstabin1 is part of the RyR1 complex in skeletal muscle and calstabin2 (FKBP12.6) is part of the RyR2 complex in cardiac muscle (25). Calstabins are immunophilins (26) with peptidyl-prolyl isomerase however this enzymatic activity does not play a role in regulating RyR channels and rather they stabilize the closed state of RyRs and prevent a Ca*^2+^* leak via the channel (18, 27).

RyR belongs to a small family of large intracellular calcium release channels, the only other member being the inositol 1,4,5-triphosphate receptor (IP_3_R) (28–30). RyR may have evolved from IP_3_R-B, which encoded an IP_3_R-like channel that could not bind IP_3_ and was replaced by RyR at the Holozoa clade (31). Invertebrates have one gene for each of RyR and IP_3_R, while vertebrates have three (RyR1-3 and IP_3_R1-3). RyRs and IP3Rs are intracellular Ca^2+^ release channels on the SR/ER and are tetramers that along with associated proteins comprise the largest known ion channel macromolecular complexes (18, 32). Defects in Ca^2+^ signaling linked to stressed-induced remodeling that results in leaky RyR channels have been implicated in heart failure (33, 34), cardiac arrhythmias (33, 35–38), diabetes (39), muscle weakness (40–43), and neurodegenerative disorders (41, 44, 45).

RyR has evolved unique SPRY domains (12, 46) that are absent in IP_3_R, one of which (SPRY2) allows RyR1 to directly interact with the L-Type Calcium channel (Cav1.1) in skeletal muscle (47). This interaction couples excitation of the sarcolemma to muscle contraction and eliminates dependence on extracellular Ca^2+^. RyR1 is remarkably well-conserved, suggesting that independence from extracellular Ca^2+^ evolved to support locomotion in higher organisms.

*UNC-68* is the RyR gene homolog in the *C. elegans* genome (48). Worms lacking both exon 1.1 and promoter1 (49), and *UNC-68 (e540)* null mutants exhibit severely defective swimming behavior and locomotive characterized by the “unc”, or “uncoordinated” phenotype (50). Treatment with ryanodine, a chemical ligand of RyR, induces contractile paralysis in wild-type *C. elegans*, whereas *UNC-68 (e540)* null mutants are unaffected by ryanodine (48, 50, 51). Ca^2+^ transients triggered by action potentials in *C. elegans* body wall muscles require *UNC-68*.

We previously reported that in aged mice (2 years old equivalent to ~80 year old humans) oxidized, leaky RyR channels contribute to loss of muscle function and impaired muscle strength (16). In the present study, we show that *UNC-68* is comprised of a macromolecular complex that is remarkably conserved compared to RyR1 and includes the channel-stabilizing subunit, *FKB-2*. Like calstabin, *FKB-2* regulates *UNC-68* by directly associating with the channel. Similar to what we previously observed in mice (15), we found age-dependent oxidation of *UNC-68*, depletion of *FKB-2* from the *UNC-68* channel complex, and reduced Ca^2+^ transients in aged nematodes. This aging phenotype was accelerated in *FKB-2 (ok3007)* worms, an *FKB-2* deletion mutant that results in leaky *UNC-68*. Competing *FKB-2* from *UNC-68* with rapamycin or FK506 (52) resulted in reduced body wall muscle Ca^2+^ transients and oxidation of *UNC-68*. Conversely, pharmacological and genetic oxidation of *UNC-68* with the reactive oxygen species (ROS)-generating drug paraquat (53) reduced *FKB-2* association with the channel and reduced contraction-associated Ca^2+^ transients. Re-associating *FKB-2* with *UNC-68* using the RyR-stabilizing drug S107 improved Ca^2+^ transients in aged nematodes. Our study provides an underlying mechanism for age-dependent loss of muscle function in *C. elegans*, progressive oxidation of *UNC-68*, which renders the channel leaky within 2 weeks compared to 2 years in mice and 80 years in humans.

## RESULTS

### Conserved evolution and architecture of UNC-68

Phylogenic analysis of RyR and FKBP among species reveals remarkable evolutionary conservation (**Figure 1A-B**). *UNC-68*, the *C. elegans* intracellular calcium release channel, shares ~40% homology with the human RyR1 (**Figure 1C**). *C. elegans FKB-2* has ~60% sequence identity with the skeletal muscle isoform calstabin1 (FKBP12) (**Figure 1D**). Based on these observations, we hypothesized that in *C. elegans, UNC-68* comprises a macromolecular complex, similar to that of mammalian RyRs. To test this hypothesis, lysates were prepared from populations of freeze-cracked wild type *C. elegans* and *UNC-68* was immunoprecipitated using mammalian anti-RyR antibody (5029) as previously described (54). The immunoprecipitates were immunoblotted to detect *UNC-68,* as well as other components of the RyR macromolecular complex including the catalytic subunit of protein kinase A (PKA_cat_), protein phosphatase 1 (PP1), *FKB-2*, and phosphodiesterase 4 (PDE-4) using mammalian anti-RyR, anti-PKA, anti-PP1, anti-calstabin, and anti-PDE-4 antibodies, respectively (**Figure 1E**). The previously published *C. elegans* anti-PDE-4 (55) was used to detect PDE-4 on the channel. Our data show that *UNC-68* comprises a macromolecular complex, similar to that found in the mammalian muscle, that includes PKA_cat_, PP1, PDE-4 and FKB-2. *UNC-68* was depleted of FKB-2 in the FKB-2 (ok3007) null mutant (**Figure 1E and G**). In the FKB-2 null *C. elegans*, *UNC-68* and the rest of the macromolecular complex could not be immunoprecipitated using an anti-FKBP antibody (**Figure 1F and H**). Taken together these data indicate remarkable evolutionary conservation of the RyR macromolecular complex.

**Figure 1:**
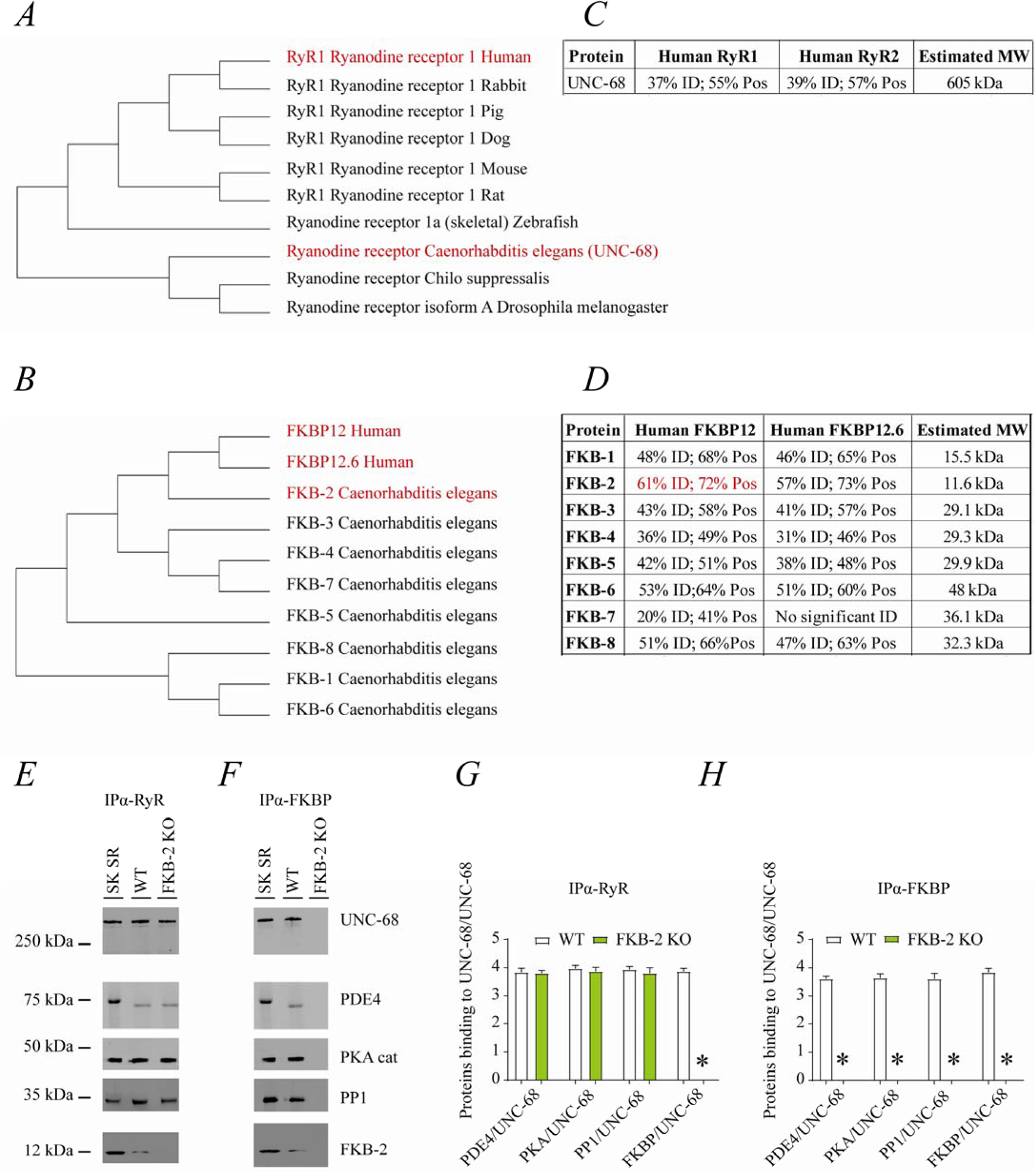
UNC-68 comprises a macromolecular complex comparable to its mammalian homologue RyR. RyR (A) and FKBP (B) evolution among species was inferred by the maximum likelihood method based on the JTT matrix-based model. (C) Homology comparison between *UNC-68* and the two human RyR isoforms (RyR1 and RyR2). (D) Homology comparison between the different FKB isoforms (1 to 8) and the human FKBP isoforms (FKBP12 and FKB12.6). UNC-68 (E) and FKB-2 (F), respectively, were immunoprecipitated and immunoblotted using anti-RyR, anti-phosphodiesterase 4 (PDE4), anti-protein kinase A (catalytic subunit; PKAcat), anti-protein phosphatase 1 (PP1), and anti-Calstabin (FKBP) antibodies in murine skeletal sarcoplasmic reticulum preparations (Sk SR), wild type populations of *C. elegans* (WT), and populations of *FKB-2 (ok3007)*. iImages show representative immunoblots from triplicate experiments. (G and H) quantification of bands intensity shown in E and F. N=6 per group. Data are means ±SEM. One-way ANOVA shows * P<0.05 WT vs. FKB-2 KO. SK SR; Sarcoplasmic reticulum fraction from mouse skeletal muscle.

### Age-dependent biochemical and functional remodeling of UNC-68

RyR1 channels are oxidized, leaky, and Ca^2+^ transients are reduced in aged mammalian skeletal muscle (15). These changes occur by two years of age in mice (15) and by 80 years of age in humans. Similarly, *C. elegans* exhibited an age-dependent decline in body wall muscle peak Ca^2+^ transients from day 3 to day 15 post-hatching (**Figure 2A-B**).

**Figure 2:**
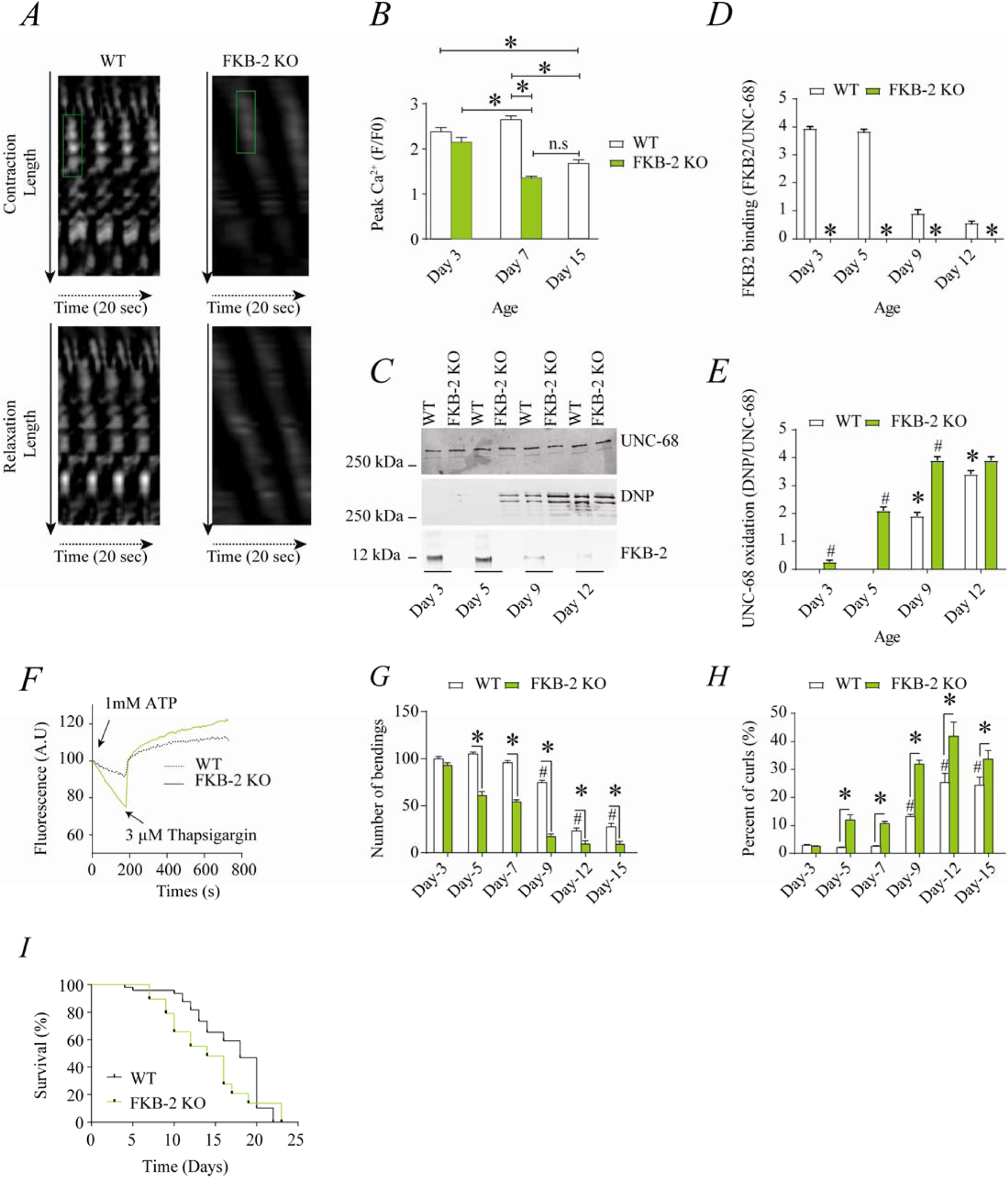
Remodeling of UNC-68 and age-dependent reduction in Ca^2+^ transients is accelerated in FKB-2 *(ok3007)* (A) Representative trace of Ca^2+^ transients from GCaMP2 WT and FKB-2 KO. Green box denotes peak fluorescence from worm’s muscle during contraction. (B) Ca^2+^ transients in age-synchronized populations of wild type and FKB-2 *(ok3007)* nematodes; (C) UNC-68 was immunoprecipitated from age-synchronized populations of mutant (FKB-2 KO) and WT nematodes and immunoblotted using anti-RyR, anti-Calstabin and DNP (marker of oxidation) antibodies. (D) and (E) Quantification of the average band intensity from triplicate experiments: band intensity was defined as the ratio of each complex member’s expression over its corresponding /UNC-68’s expression Data are means ±SEM. * P<0.05 WT vs. FKB-2 KO in panel D, # P<0.05 WT vs. FKB-2 KO in panel E, * P<0.05 WT at Day3 vs. WT at Day5 and Day9. (F) Ca^2+^ leak assay performed with microsomes from WT and FKB-2 KO worms (Day 5). Ca^2+^ uptake into the microsomes was initiated by adding 1mM of ATP. Then, 3µM of thapsigargin was added to block the SERCA activity. Increased fluorescence is proportional to the spontaneous Ca^2+^ leakage throughout *UNC-68.* (G) Graph showing number of bends recorded for WT vs FKB-2 KO worms at six distinct ages (Day 3, 5, 7, 9, 12, 15). (H) Number of curling events were calculated as a percentage of the overall motility (curls/bends). N = ~60 worms per group, with the exception of Day 15 (as fewer worms were alive at this time point). Day 15 = ~40 worms. (I) percentage of survival of WT and FKB-2 KO worms. Data are means ±SEM. One Way ANOVA shows * P<0.05 WT vs. FKB-2 KO, # P<0.05 WT at Day3 vs. Day5-7-9-12 and 15.

RyR1 oxidation has been linked to SR Ca^2+^ leak and impaired muscle function during extreme exercise and in heart failure and muscular dystrophies (23, 56, 57). Furthermore, we have previously reported that oxidation of RyR1 and the subsequent intracellular Ca^2+^ leak are underlying mechanisms of age-related loss of skeletal muscle specific force (force normalized to the cross sectional area of muscle) (15). WT *UNC-68* was depleted of *FKB-2* (**Figure 2C-D**) and oxidized (**Figure 2C-E**) in an age-dependent manner. These changes mirror those occurring with extreme exercise in mice and humans (23) and in a murine model of Duchenne Muscular Dystrophy (mdx mice) characterized by impaired muscle function (56). Importantly, by 80 years of age, ~50% of humans develop severe muscle weakness that is a strong predictor of mortality due to falls, gait imbalance, and related factors (58).

### FKB-2 deficiency accelerates age-dependent reduction in Ca^2+^ transient that drives muscle contraction

*FKB-2* deficiency accelerated age-dependent reduction in body wall muscle Ca^2+^ transients (**Figure 2A-B**), with peak Ca^2+^ in *FKB-2 (ok3007)* worms at day 7 being significantly lower than that of wild type at the same age. Similarly, *UNC-68* was significantly more oxidized (day 3 to 9) in *FKB-2 (ok3007)* worms compared to wild type (**Figure 2C, D, E**).

### UNC-68 channels are leaky in the absence of FKB-2

To further demonstrate that *UNC-68* channels lacking *FKB-2* are inherently ‘leaky’, we used an assay that can monitor the rate of Ca^2+^ released from the SR. Synchronized worm microsomes (day 5) were mixed with the Ca^2+^ dye Fluo-4 and baseline fluorescent measurements were taken before adding 1mM of ATP. By activating the sarco/endoplasmic calcium ATPase (SERCA) with ATP, cytosolic Ca^2+^ is pumped into the microsomes until the cellular compartment reaches capacity, resulting in a subsequent decrease in Fluo-4 fluorescence. Once the fluorescence level plateaus, thapsigargin (SERCA antagonist) is added to block Ca^2+^ re-uptake into the SR. The rate at which the fluorescence increases directly correlates with the amount of Ca^2+^ entering the cytoplasm: a higher increase of cytosolic Ca^2+^ compared to WT controls is suggestive of leaky *UNC-68* channels. Our data show that *UNC-68* from FKB-2 KO worms had a higher rate of SR Ca^2+^ leak following thapsigargin administration compared to the WT channels (**Figure 2F**). This is corroborated by our previous findings, where disruption of RyR-calstabin binding increases the SR Ca^2+^ leak in mammalian tissues (16, 59).

### UNC-68 channel leak impairs exercise capacity

In mammals, calstabin regulation of RyR is tightly coupled to beta-adrenergic signaling (60), and it is known that calstabin KO mice must undergo exercise stress before demonstrating a distinct muscle phenotype (23). Our method of inducing exercise stress in the worm was to place it in M9 buffer and observe it swimming, a well-described behavioral assay (61). By using an extended time trial of 2 hours, the worms fatigue and exhibit exercise-induced stress similar to that observed in mammals. Our data show a defect in *FKB-2 KO* swimming behavior over the course of its lifespan when compared to the WT. *FKB-2 KO* worms had decreased bending activity earlier in life, beginning at day 5, and an increased proportion of curling, a sign of fatigue (**Figure 2G-H**). Throughout midlife, the *FKB-2* KO worms lag significantly behind their age-matched WT counterparts, suggestive of decreased muscle function. Furthermore, *FKB-2* KO worms exhibit reduced life span compared to WT (**Figure 2I**).

### Pharmacologically mimicking aging phenotype affects Ca^2+^transient and impairs exercise capacity

*FKB-2* was competed off from the *UNC-68* macromolecular complex using rapamycin or FK506 (**Figure 3**). Both rapamycin and FK-506 bind to calstabin and compete it off from RyR channels, resulting in leaky channels and release of SR Ca^2+^ in the resting state (27, 62, 63). Rapamycin-FKBP12 inhibits mTOR and FK506 inhibits calcineurin, indicating that these distinct non-*UNC-68* related actions do not account for the observed SR Ca^2+^ leak.

**Figure 3:**
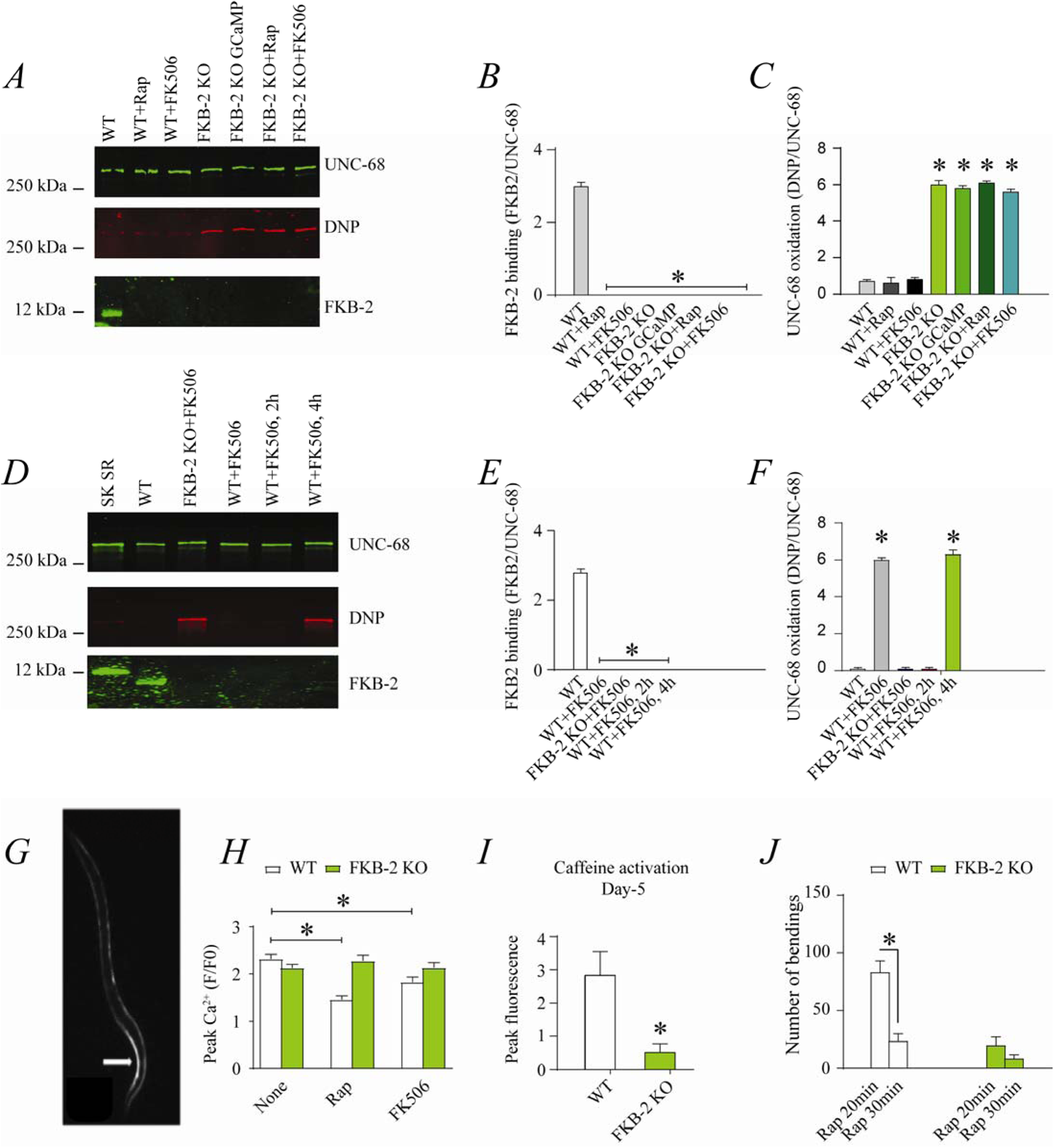
Depleting FKB-2 from UNC-68 causes UNC-68 oxidation. (A) UNC-68 was immunoprecipitated and immunoblotted using anti-RyR, anti-Calstabin and DNP (marker of oxidation) antibodies in nematodes acutely treated with 15μM and 50μM rapamycin and FK506, respectively. (B-C) Quantification of the band intensity shown in (A): band intensity was defined as the ratio of either DNP (marker of *UNC-68* oxidation) or FKB-2 binding over its corresponding /UNC-68’s expression. (D) UNC-68 was immunoprecipitated after 2hrs, and 4 hrs FK506 exposure. Representative immunoblots from triplicate experiments. (E-F) Quantification of the band intensity shown in (D): band intensity was defined as the ratio of either DNP (marker of *UNC-68* oxidation) or FKB-2 binding over its corresponding /UNC-68’s expression. G) Representative image of caffeine activated calcium transient in GCaMP2 WT; arrow denotes peak fluorescence in body-wall muscle. (H) Ca^2+^ transients in age-synchronized populations of wild type and FKB-2 (ok3007) nematodes treated with 15μM and 50μM rapamycin and FK506, respectively. (I) Fluorescence intensity following caffeine activation in age-matched GCaMP2: WT versus GCaMP2: FKB-2 KO worms at Day 5. (J) Graph showing number of bends recorded for WT vs FKB-2 KO worms (Day 5) treated for 20 and 30 minutes with 15μM and 50μM rapamycin and FK506, respectively. N≥15 per group. Data are means ±SEM. One-way ANOVA shows * P<0.05 vs. WT for results shown in panel A to F. Two-way ANOVA was used for results comparison in panel H and t-test was used for results shown in I and J. Sarcoplasmic reticulum fraction from mouse skeletal muscle.

Age-synchronized young *C. elegans* were treated with rapamycin or FK506. Acute treatment with FK506 or rapamycin each independently caused depletion of FKB-2 from the channel (**Figure 3A, B, C**). Furthermore, longer treatment of WT worms with FK506 caused oxidation of *UNC-68*, demonstrating a relationship between depletion of *FKB-2* and oxidation of *UNC-68* (**Figure 3D, E, F**). Ca^2+^ transients were measured in partially immobilized transgenic nematodes expressing the genetically encoded Ca^2+^ indicator, P*myo-3*::GCaMP2, in the body wall muscle cells (64, 65) (**Figure 3G**). Pharmacologic depletion of *FKB-2* from *UNC-68* by rapamycin or FK506 treatment caused reduced body wall muscle Ca^2+^ transients in wild type *C. elegans* (**Figure 3H**). When FKB-2 was genetically depleted from the *UNC-68* complex, as in the *FKB-2 (ok3007)* nematodes, treatment with rapamycin or FK506 had no effect on the Ca^2+^ transients (**Figure 3H**).

Continuous Ca^2+^ leak via *UNC-68* would be expected to result in depleted SR Ca^2+^ stores; therefore, we utilized a common technique from the mammalian RyR literature to evaluate the SR Ca^2+^ stores. In brief, an activating concentration of caffeine is used to fully open the RyR channel, leading to a rapid release of calcium from the SR into the cytoplasm. This increase can be approximated using a previously targeted, florescent calcium dye or indicator. Caffeine was applied to day 5 cut worms (**Figure 3I**) and the amount of fluorescence given off by GCaMP2 was measured. The GCaMP2-WT worms demonstrated a strong Ca^2+^ transient within 10 seconds of caffeine administration, while GCaMP2-*FKB-2* KO worms failed to produce a response, suggesting that their SR Ca^2+^ stores were too low to elicit one. Interestingly, GCaMP2-KO worms were observed as having very high background fluorescence, which may indicate an increase in cytosolic Ca^2+^ from passive *UNC-68* leak. Indeed, rapamycin altered swimming behavior of WT but not *FKB-2* KO worms in a time dependent manner (**Figure 3J**). Taken together with our Ca^2+^ transient data, the observed muscle phenotype appears to be the result of *UNC-68* channel leak. These data suggest that rendering *UNC-68* channels leaky by removing *FKB-2* depletes SR Ca^2+^, resulting in reduced Ca^2+^ transients and weakened muscle contraction.

### Oxidation of UNC-68 causes reduced body wall muscle Ca^2+^ transients

To investigate the individual effect of age-dependent *UNC-68* oxidation independent of the other confounding variables involved in aging (4), we introduced a pharmacological intervention mimicking the aged state in young adult nematodes. Treating young adult nematodes with the superoxide-generating agent paraquat (53) increased oxidation of *UNC-68* and depletion of *FKB-2* from the channel in a concentration-dependent manner (**Figure 4A, B, C**). Furthermore, contraction-associated Ca^2+^ transients decreased with paraquat treatment in a concentration-dependent manner (**Figure 4D**). Indeed, treatment with antioxidant N-Acetyl-L-cysteine (NAC) improved Ca^2+^ transient in *FKB-2* KO worms (**Figure 4E**). These data indicate that both *UNC-68* oxidation and *FKB-2* depletion independently contribute to the observed aging body wall muscle deterioration.

**Figure 4:**
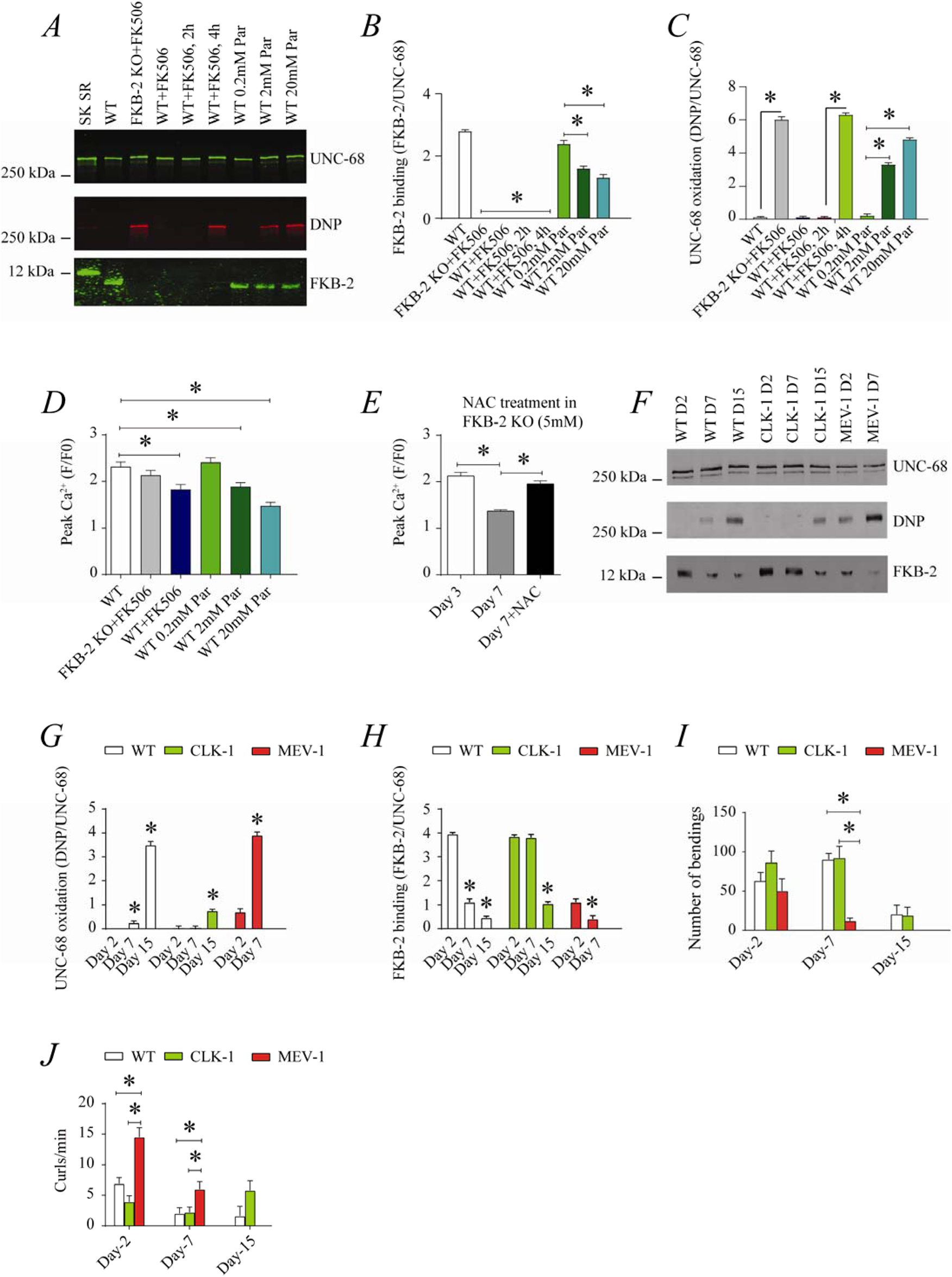
UNC-68 oxidation causes defective Ca^2+^ handling; (A) UNC-68 was immunoprecipitated and immunoblotted using anti-RyR, anti-Calstabin and DNP (marker of oxidation) antibodies in nematodes acutely treated for 15-20 minutes with FK506 or paraquat at increasing concentration. (B-C) Quantification of the band intensity shown in (A): band intensity was defined as the ratio of either DNP (marker of *UNC-68* oxidation) or FKB-2 binding over its corresponding/UNC-68’s expression. (D) Contraction-associated Ca^2+^ transients measured in young age-synchronized WT nematodes treated for 15-20 minutes with increasing concentration of paraquat. (E) Contraction-associated Ca^2+^ transients measured in FKB-2 KO nematodes treated with the antioxidant N-acetylcysteine (NAC) at 5mM. (F) *UNC-68* was immunoprecipitated and immunoblotted using anti-RyR, anti-Calstabin and DNP (marker of oxidation) antibodies in WT, the long lived (CLK-1) and the short lived (MEV-1) nematodes at day 2, 7 and 15. (G-H) Quantification of the average band intensity from triplicate experiments: band intensity was defined as the ratio of each complex member’s expression over its corresponding /UNC-68’s expression. (I) Graph showing number of bends recorded for WT vs CLK-1 and MEV-1 worms at three distinct ages (Day 2, 7, 15). J) Number of curling events were calculated as a percentage of the overall motility (curls/bends). N≥.20 per group. Data are means ±SEM. One-way ANOVA shows * P<0.05. Two-way ANOVA was used in panel I and J. Sarcoplasmic reticulum fraction from mouse skeletal muscle.

To better clarify the role of oxidative stress in age-dependent *UNC-68* remodeling and Ca^2+^ leak, we used two mutant mitochondrial electron transport chain (ETC): the complex I mutant, CLK-1, and the complex II mutant, MEV-1. CLK-1 worms contain a Complex I-associated mutation where they cannot synthesize their own ubiquinone (UQ), a redox active lipid that accepts and transfers electrons from Complex I or II to Complex III in the ETC. The reduction in Complex I activity of CLK-1 is associated in long living worms (66–68). In contrast, MEV-1 worms contain a complex II (succinate dehydrogenase) cytochrome B560 mutation (69–71), preventing electron transfer from succinate to fumarate and caused mitochondrial ROS production, which is associated with decreased lifespan, averaging only 9 days (70).

Interestingly, we have seen increased *UNC-68* oxidation and *FKB-2* depletion in the short-lived mutant (MEV-1) compared to WT and long-lived mutant (CLK-1) worms (**Figure 4F, G and H**). Indeed, MEV-1 worms exhibited reduced exercise capacity compared to WT and CLK-1 worms (**Figure 4I-J**).

### UNC-68 Ca^2+^ channel is a potential therapeutic target in aging

The small molecule Rycal S107 inhibits SR Ca^2+^ leak by reducing the stress-induced depletion of calstabin from the RyR channel complex (56, 72). Here, we show that treatment with S107 (10μM) for 3-5 hours re-associated *FKB-2* with *UNC-68* without significant effect on the channel oxidation (**Figure 5A, B, C**). Furthermore, treatment with S107 improved peak Ca^2+^ in an *FKB-2* dependent manner, as demonstrated by the fact that treating the *FKB-2* KO worms results in no change in peak Ca^2+^ (**Figure 5D-E**). Interestingly, S107 treatment reduced age-dependent impairment of exercise capacity in WT worms at day 15 (**Figure 5F**). Of note, S107 has no effect on the WT worms life span (**Figure 5G**). Furthermore, the treatment of the short-lived worms, MEV1, with S107 restored the FKB-2 association with *UNC-68*, despite the persistence of the channel oxidant (**Figure 5H, I and J**).

**Figure 5:**
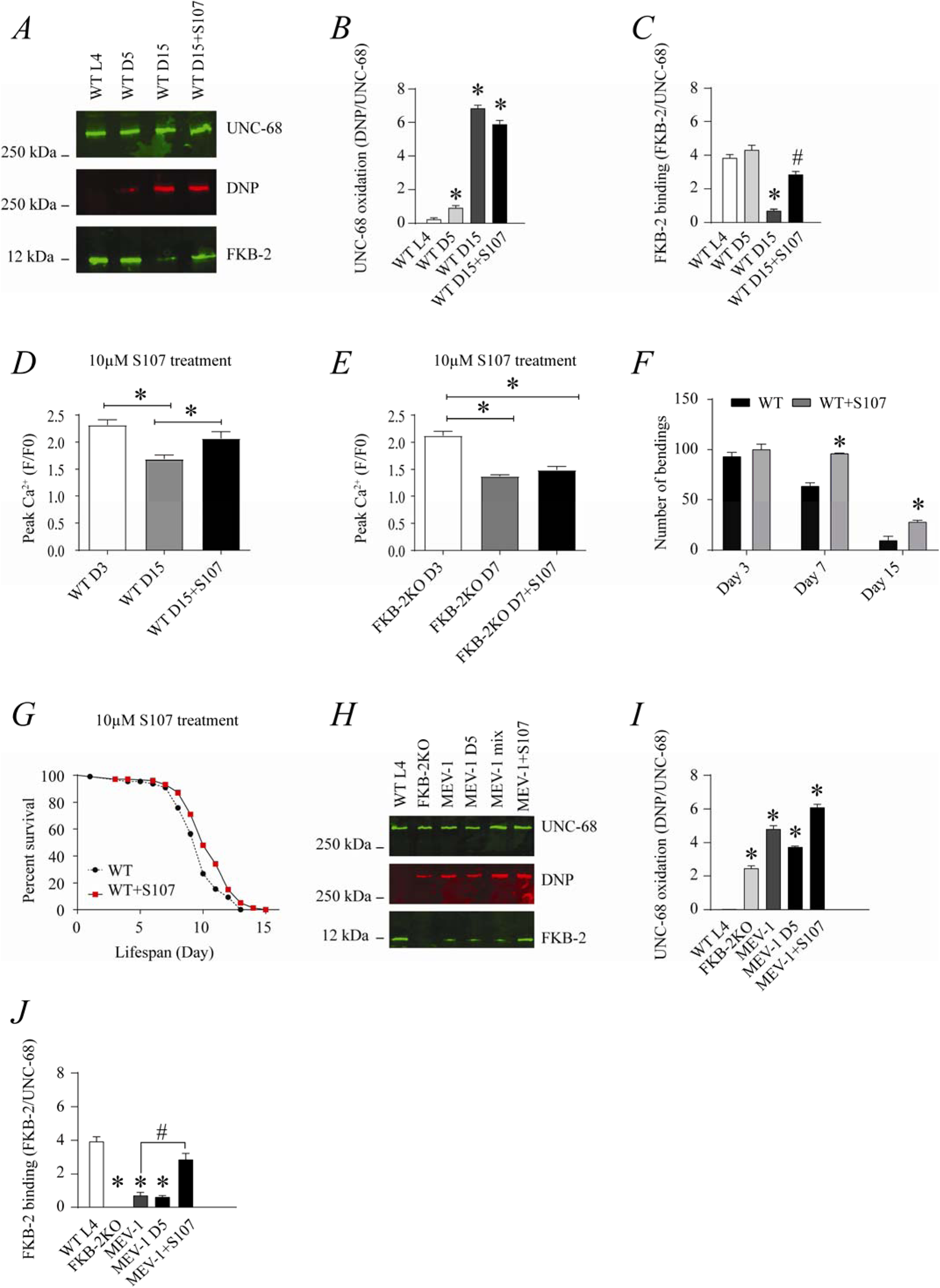
The RyR-stabilizing drug S107 increases body wall muscle Ca^2+^ transients in aged *C. elegans*; (A) UNC-68 was immunoprecipitated and immunoblotted with anti-RyR, anti-Calstabin and DNP (marker of oxidation) in aged nematodes with 10µM of S107 (5hrs). (B-C) Quantification of the band intensity shown in (A): band intensity was defined as the ratio of either DNP (marker of *UNC-68* oxidation) or FKB-2 binding over its corresponding /UNC-68’s expression. Data are means ±SEM. * P<0.05 vs. WTL4, # P<0.05 WT-D 15 vs. WT D15+S107. (D-E) Contraction-associated Ca^2+^ transients were measured in age-synchronized WT (D) and (E) *fkb-2 (ok3007)* treated with S107 as indicated. (F) Graph showing number of bends recorded for WT vs WT treated with S107 worms at different age (Day 3, 7, 15 days). (G) Percent of survival of WT vs WT treated with S107 nematodes. (H) UNC-68 was immunoprecipitated and immunoblotted with anti-RyR, anti-Calstabin and DNP (marker of oxidation) in short-lived nematodes (MEV-1) with S107 treatment (3-5hrs). (I-J) Quantification of the band intensity shown in (H): band intensity was defined as the ratio of either DNP (marker of *UNC-68* oxidation) or FKB-2 binding over its corresponding/UNC-68’s expression. N≥20 per group. Data are means ±SEM. One-way ANOVA shows * P<0.05 vs. In panel F, a t-test was used to compare WT and WT+S107 for each day. WTL4 unless otherwise indicated. #P<0.05 MEV-1, vs MEV-1+S107.

## DISCUSSION

Taken together, our data show that the *C. elegans* intracellular Ca^2+^ release channel *UNC-68* comprises a macromolecular complex which is highly conserved throughout evolution from nematodes to humans. In nematodes, the *UNC-68* macromolecular complex is comprised of a similar array of regulatory subunits as the mammalian RyR1 channels: a phosphodiesterase PDE-4, a protein kinase PKA, a protein phosphatase PP1, and the immunophilin, *FKB-2*. Binding of *FKB-2* (the *C. elegans* homolog of the mammalian RyR stabilizing protein calstabin) to the *UNC-68* channel is required to prevent a pathological leak of intracellular Ca^2+^, similar to the manner observed in mammalian muscle (15). Aged *C. elegans* exhibit reduced Ca^2+^ transients, as well as oxidized *UNC-68 channels* and depleted *FKB-2* by ~2 weeks of age. Each of these post-translational modifications of *UNC-68* likely contribute to the aging phenotype, as pharmacologically depleting *UNC-68* of *FKB-2* and oxidizing the channel in young nematodes each independently result in a premature aging phenotype and reduce contraction-associated Ca^2+^ transients. Genetic *FKB-2* deficiency causes an accelerated aging phenotype; Ca^2+^ transients are reduced in younger populations of *FKB-2 (ok3007)* nematodes and *UNC-68* is oxidized at an earlier time point in these mutants relative to wild type. Treating aged wild type nematodes with the RyR-stabilizing drug, S107, re-associates *FKB-2* with *UNC-68* and increases the Ca^2+^ transients, indicating that *UNC-68* dysfunction associated with oxidation-induced remodeling of the *UNC-68* macromolecular complex is likely an underlying mechanism of age-dependent decrease in Ca^2+^ transients in function *C. elegans* body wall muscle.

*C. elegans* exhibit an aging muscle phenotype similar to age-dependent loss of muscle function in humans (15). This is characterized by impaired locomotion, reduction in muscle cell size associated with loss of cytoplasm and myofibrils, and progressive myofibril disorganization (4). However, specific body wall muscle proteins involved in the *C. elegans* aging phenotype have not been determined. Here, we show that *UNC-68* is oxidized in aged nematodes and depleted of the channel-stabilizing protein, *FKB-2*. Our group has reported similar remodeling of RyR1 in skeletal muscle from aged mice (15) and in murine models of Muscular Dystrophies (56), all of which exhibit intracellular Ca^2+^ leak and reduced muscle specific force production.

Though the oxidative stress theory of aging was first proposed in 1956 (73, 74), there is still substantial controversy surrounding the role of ROS in aging. For example, deletion or overexpression of the ROS detoxification enzyme superoxide dismutase (SOD) has little effect on life span in *C. elegans* (75, 76). However, loss of *sesn-1*, the gene encoding sestrin, an evolutionarily conserved protein required for regenerating hyper-oxidized forms of peroxiredoxins and for ROS clearance, causes reduced lifespan (77). Furthermore, ROS levels measured *in vivo* in *C. elegans* increase with age (78). Other oxidative/antioxidative genes are involved in ROS production and may play a crucial role in the *UNC-68* oxidation (**Table S1**).

While the Free Radical Theory of Aging (FRTA) has taken a hit due to multiple observations that contradict the notion of a link between reduced oxidative load and longevity the preponderance of data shows a correlation between oxidative damage and reduced lifespan (79). Moreover, there is no doubt that reduced muscle function is detrimental to survival (80). The present study shows that a key effector of age-dependent oxidative overload, RyR1 channel leak and the resulting muscle dysfunction, occur approximately 2,000 times faster in *C. elegans* compared to *Homo sapiens* and 50 times faster than in *Mus musculus.* Since the target system, RyR1/UNC68, are remarkably conserved and underlay dramatically similar physiological functions (namely SR Ca^2+^ release required for muscle contraction) the cause for the accelerated kinetics of aging must be determined elsewhere and in an unrelentingly constant manner as exemplified by the rigid control of species lifespan. There is however, only one known case of a significant prolongation of lifespan in a species: *Homo sapiens*. Indeed, the average lifespan in the U.S. has doubled in the past century (81) largely due to improved sanitation and related public health measures that protect against communicable diseases, the present pandemic notwithstanding. This suggests that both environmental an d intrinsic biological constraints can determine the limits of lifespan. Since we are a species that can remodel our environment to a greater extent than any other we have been able to double our lifespan by improving the environment although now global warming threatens to reverse this achievement. The unanswered question remains what are the intrinsic biological constraints on a given species longevity?

It would be interesting to know if the increased *UNC-68* oxidation and subsequent reduction in body wall muscle Ca^2+^ transients are a result of globally increased ROS levels or increased ROS levels in *UNC-68*-surrounding microdomains. For example, we have previously shown that inducing RyR leak in enzymatically dissociated skeletal muscle cells causes increased mitochondrial membrane potential and mitochondrial ROS production (15). Based on these data, we have proposed a model in which RyR1 leak (due to age-dependent oxidation of the channel) causes mitochondrial Ca^2+^ overload, resulting in ROS production, thus leading to further oxidation of RyR1 and exacerbation of the SR Ca^2+^ leak. This creates a vicious cycle between RyR1 and mitochondria that contributes to age-dependent loss of muscle function (82).

We also demonstrate that the putative null mutant, *FKB-2 (ok3007),* prevents *FKB-2* from co-immunoprecipitating with *UNC-68*. The aging phenotype that we characterize in wild type nematodes (biochemically modified *UNC-68* and reduced Ca^2+^ transients) is accelerated in *FKB-2 (ok3007).* There are eight *FKBs* that are homologous to mammalian calstabin in the *C. elegans* genome; *FKB-1* and *FKB-8* both have ~50% sequence identity to calstabin. Further studies could elucidate the possibility that in the absence of *FKB-2*, another *FKB* may stabilize *UNC-68*, in particular the aforementioned *FKB-8* (its gene is in close proximity to that of *FKB-2* on chromosome 2) and *FKB-1* (most similar to FKB-2 in terms of molecular weight). However, this appears not to be the case, since the aging phenotype is accelerated in the absence of *FKB-2*. Another key question is why *UNC-68* becomes oxidized within two weeks, whereas the same post-translational modification requires two years in mice and 80 years in humans (5). Given the high degree of conservation of RyR and other members of the complex (**Figure 1**), it is feasible that genetic screens in organisms such as *C. elegans* and *Drosophila* will yield additional crucial mediators that are common among species and explain disparities in lifespan such as genes-genes interactions, epigenetics or architecture and gating of key proteins involved in aging such as RyR. Indeed, despite the conservative evolution of *UNC-68*, the channel contains higher number of methionine (3.5%) and serine (7.2%) (**Table S2**) compared to the human RyR1 (2.9% and 5.9% respectively). Methionine are a primary target of oxidative stress that might cause defects in the channel gating and alter Ca^2+^ release. Disparities in RyR1 serine residues among species, which are phosphorylation by protein kinases in response to stress, can cause conformational changes to the channel, exposing thereby more residues to oxidation and could be a potential mechanism responsible of the accelerated RyR1 oxidation in *C. elegans*.

Taken together, our data indicate that the *C. elegans* homolog of RyR, *UNC-68*, is comprised of a macromolecular complex and regulated by the immunophilin, *FKB-2*. We have identified age-dependent reduction in body wall muscle Ca^2+^ transients in nematodes that is coupled to oxidation and remodeling of *UNC-68*. SR Ca^2+^ stores are depleted in *FKB2*-KO worms, suggesting passive *UNC-68* leak. This observation is supported by the Ca^2+^ leak assay results, which show that *FKB-2* regulation is critical in preventing *UNC-68* channels from aberrantly ‘leaking’ Ca^2+^ into the cytoplasm. With reduced SR Ca^2+^, *UNC-68* fails to release the burst of Ca^2+^ required for normal excitation-contraction coupling, leading to impaired muscle function. Loss of muscle function is evident in the *FKB-2* KO worms during swimming trials, as middle-aged worms performed worse than their age-matched WT controls. Furthermore, our data strongly suggest a role for *FKB-2* and *UNC-68* in the age-dependent changes in Ca^2+^ signaling, as treatment with the pharmacological RyR stabilizer S107 increases body wall muscle Ca^2+^ transients.

## METHODS

### C. elegans strains and culture conditions

Worms were grown and maintained on standard nematode growth medium (NGM) plates on a layer of OP50 *Escherichia coli* at 20°C, as described (50). N2 (Bristol) and *fkb-2 (ok3007)* were provided by the Caenorhabditis Genetics Center (University of Minnesota). *fkb-2 (ok3007)* was backcrossed six times. The transgenic strain expressing P*myo-3*:GCaMP2 was kindly provided by Zhao-Wen Wang, University of Connecticut Health Center (65). P*myo-3*: GCaMP2 was subsequently crossed into *fkb-2 (ok3007)* for measurement of contraction-associated Ca^2+^ transients.

### Age synchronization

Adult worms at the egg-laying stage were treated with alkaline hypochlorite solution to obtain age-synchronized populations and eggs were plated on NGM plates, as described (83). For experiments requiring aged worms, age-synchronized animals at the L4 stage were collected in M9 buffer and plated on NGM plates containing 5-fluoro-2’-deoxyuridine (FUDR, Sigma, 50 μM) to prevent egg-laying (84).

### Immunoprecipitation and immunoblotting

Nematodes were grown under standard conditions. For protein biochemistry experiments, a procedure to crack nematodes in a solubilizing and denaturing buffer was adapted (85). Briefly, worms were washed and collected with M9 buffer, centrifuged for 2 minutes at 1,000 rpm three times to wash. Worms were allowed to settle to the bottom of the collection tube by sitting on ice for ~5 minutes. Fluid was removed and the worm pellet was snap frozen in liquid nitrogen. Frozen pellets containing whole nematodes were rapidly thawed under warm running water. A volume of Nematode solubilization buffer equal to the volume of the worm pellet was added (Nematode solubilization buffer: 0.3% Ethanolamine, 2 mM EDTA, 1 mM PMSF in DMSO, 5 mM DTT, 1x protease inhibitor) and tubes were microwaved (25 s for 100 μl pellet; time was increased for greater volumes). Lysates were then quickly drawn into a syringe through a 26-gauge needle and forced back through the needle into a new collection tube on ice. Samples were centrifuged at 1,000 rpm for 2 min to remove insoluble material and the supernatant was transferred to a new tube on ice. Lysates were snap frozen and stored in at −80°C.

A anti-mammalian RyR antibody (4 μg 5029 Ab (86)) was used to immunoprecipitate *UNC-68* from 100 μg of nematode homogenate. Samples were incubated with the antibody in 0.5 ml of a modified RIPA buffer (50 mM Tris-HCl pH 7.4, 0.9% NaCl, 5.0 mM NaF, 1.0 mM Na3VO4, 1% Triton-X100, and protease inhibitors) for 1 hr at 4°C. The immune complexes were incubated with protein A Sepharose beads (Sigma, St. Louis, MS) at 4°C for 1 hour, after which time the beads were washed three times with buffer. Proteins were size-fractionated by SDS-PAGE (6% for *UNC-68*, 15% for FKB-2) and transferred onto nitrocellulose membranes for 1 hour at 200 mA (SemiDry transfer blot, Bio-Rad). After incubation with blocking solution (LICOR Biosciences, Lincoln NE) to prevent non-specific antibody binding, immunoblots were developed using antibodies against RyR (5029, 1:5000), PKAcat (Santa Cruz Biotechnology, sc-903, 1:1000), PDE4 (kindly provided to us by Kenneth Miller, Oklahoma Medical Research Foundation, Oklahoma City, Oklahoma), PP1 (sc6104, 1:1000) or an anti-calstabin antibody (Santa Cruz 1: 2,500). To determine channel oxidation, the carbonyl groups on the protein side chains were derivatized to 2,4-dinitrophenylhydrazone (DNP-hydrazone) by reaction with 2,4-dinitrophenylhydrazine (DNPH) according to manufacturers (Millipore) instructions. The DNP signal on immunoprecipitated *UNC-68* was determined by immunoblotting with an anti-DNP antibody (Millipore, 1:1000). All immunoblots were developed and quantified using the Odyssey Infrared Imaging System (LICOR Biosystems, Lincoln, NE) and infrared-labeled secondary antibodies. In addition, immunoblotting and immunoprecipitation of the *UNC-68* macromolecular complex were conducted using another anti-calstabin antibody (1:2000, Abcam) and the same methods as described.

### Imaging contraction-associated body wall muscle Ca^2+^ transients

Spontaneous changes in body wall muscle Ca^2+^ were measured in nematodes expressing GCaMP2 by fluorescence imaging using a Zeiss Axio Observer inverted microscope with an electron-multiplying CCD camera (Photometrics Evolve 512) and an LED light source (Colibri). Nematodes were partially immobilized by placing them individually into a 5-10 μl drop of M9 buffer, suspended between a glass slide and coverslip. Twenty-second videos of individual nematodes were recorded.

### Analyzing contraction-associated body wall muscle Ca^2+^ transients

Contraction-associated body wall muscle Ca^2+^ transients were analyzed using an Interactive Data Language (IDL)-based image quantification software that was developed for this purpose in our laboratory. For each twenty-second video, signals from the body wall muscles in nematodes expressing GCaMP2 fluorescence were analyzed using an edge-detection algorithm from each frame as “line-scan” images, with the nematode perimeter on the y-axis and time (s) on the x-axis (87, 88). These images were then quantified based on the average of the peak Ca^2+^ fluorescence signal on the worm muscle wall.

*Drug treatment* To pharmacologically deplete FKB-2 from *UNC-68*, nematodes were treated for 15 minutes with 15 μM rapamycin or imaging 50 μM FK506, respectively. To re-associate FKB-2 and *UNC-68*, aged nematodes were treated with 10 μM S107 for 3-5 hrs. Oxidative stress was induced in the worms using 20 mM paraquat, a known generator of superoxide (89). Nematodes were grown in standard conditions, age-synchronized as described, washed and collected with M9 buffer, then centrifuged for 2 min at 1,000 rpm three times. Worms were allowed to settle to the bottom of the collection tube by sitting on ice for ~5 min. Fluid was removed, the worm pellet was gently resuspended in M9 containing the appropriate drug concentration and gently rocked on a shaker at RT for the indicated time periods. Collection tubes were centrifuged for 2 min at 1,000 rpm and M9 containing drug was removed and replaced with M9. Biochemistry or Ca^2+^ measurements were then conducted as previously described (16).

### Measuring SR Ca^2+^ stores using caffeine activation

Age-synchronized GCaMP2: WT and GCaMP2: FKB-2 KO were grown on NGM plates at 20°C they were separated from their progeny and left undisturbed until Day 5. Individual worms were placed in a drop of M9 on a coverslip. The liquid was carefully wicked away using KIMTECH wipes until only a sliver of moisture surrounded the worm. The worm was quickly glued down to the coverslip using a tiny drop of DermaWorm applied to the head and tail of the worm before the worm desiccated. 80 μl of M9 buffer was added immediately afterwards to polymerize the glue. Once the worm was secure, a clean lateral cut to the immediate tail region was made using a 20G 1½ needle (adapted from Wang ZW et al. Neuron 2011^48^). An additional 170μl of M9 buffer was applied for a total of 250μl. The completed preparation was placed on the platform of a Zeiss confocal microscope; after 1 min at baseline, 25 mM of caffeine was added to an equal volume of M9 solution. The resulting body-wall transients were recorded for 1 min.

### Calcium leak assay

Worm microsomes from Day 5 synchronized populations (0.25 mg) were added to 150 μl solutions that contained: 20 mM HEPES ph 7.2, 7 mM NaCl,, 1.5 mM MgCl2, 0.1 mM EGTA Ca^2+^buffer (free [Ca^2+^] 0.3 μM), 8 mM phosphocreatine, and 0.02 mM fluo-4, pH 6.8. Ca^2+^ uptake was initiated by adding of 1 mM ATP, which was allowed to plateau after 3 min. After the Ca^2+^ uptake had stabilized, 3 μM thapsigargin was added to inhibit SERCA activity and the resulting Ca^2+^ leak was quantified fluoroscopically using a Tecan infinite F500 plate reader for 9 min.

### Swimming behavior

Standard M9 buffer was mixed with 2% agar and poured into 96 well plates to create a planar surface for analyzing worm swimming behavior. Once the mixture had polymerized, approximately 180 μl of M9 was pipetted on top of the agar bed and age-synchronized worms from one of two groups (WT or FKB-2 KO) were placed individually into each well. To assess differences in exercise fatigue, worms were allowed to swim freely in M9 buffer for 2 hrs; swimming bends and curls (61) were recorded by eye for 1 min.

Representative videos were taken of each group and investigators were blinded over the course of each experiment. All recordings were made in duplicate.

### Statistical analysis

All results are presented as mean ± SEM. Statistical analyses were performed using an unpaired two-tailed Student’s t test (for 2 groups) and one-way ANOVA with Tukey-Kramer test (for 3 or more groups), unless otherwise indicated. P values <0.05 were considered significant. All statistical analyses were performed with Prism 8.0.

**Table S1:** Oxidative regulators in C. elegans vs mice vs humans.

| **Protein** | ***FOXO***<br>**(Forkhead**<br>**Box Class O)** | ***CAT***<br>**(Catalase)** | ***SOD***<br>**(Superoxide**<br>**Dismutase)** | ***PRDX/PRX***<br>**(Peroxiredoxin)** | ***TRX*(Thioredoxin)** | ***GLRX***<br>**(Glutaredoxins)** |
| --- | --- | --- | --- | --- | --- | --- |
| **Function** | Transcription<br>Factor | Antioxidative<br>Enzyme | Antioxidative<br>Enzyme | Antioxidative<br>Enzyme | Antioxidative<br>Enzyme | Antioxidative<br>Enzyme |
| **Human** | FOXO1,3,4,6 | CAT | SOD1, 2, 3 | PRDX/PRX<br>1,2,3,4,5,6 | TRX | GLRX 1,2,3,5 |
| **Murine** | FOXO1,3,4,6 | cat | Sod 1, 2,3 | Prdx | Trx | Glrx 1,2,3,5 |
| ***C. elegans*** | DAF-16 | Ctl 1, 2,3 | Sod 1,2,3,4,5 | prdx 2, 3, 6 | trx 1,2,3,4,5 | glrx 3, 5, 10, 21, 22 |
| **Location**<br>**human vs**<br>***C. elegans*** | Nucleus<br>Vs<br>Nucleus | Peroxisome<br>vs<br>Peroxisome | Cytoplasm,<br>nucleus and<br>mitochondria<br>vs cytoplasm<br>and<br>mitochondria | Cytoplasm vs<br>cytoplasm | Nucleus and<br>cytoplasm vs<br>nucleus | Cytoplasm,<br>mitochondria vs<br>mitochondrial matrix |
| **Enzymatic**<br>**activity** | None | Human:<br>~50U/mg<br>(skin) and<br>98.6 MU/l<br>(blood);<br>Mouse: 4-18U/mg<br>(Liver);<br>Worm: 35-55U/mg over<br>lifespan | SOD-1<br>Human: ~30<br>U/mg (skin)<br>Mouse: 4<br>U/mg; SOD-2 | Human & Rat:<br>40<br>(mmol/min/mg) | Human:<br>0.15/100ug x 10<sup>3</sup> | Human:10nmol/min/<br>mg in lung tissues<br>Mouse : Identical to<br>human |
| **Disruption**<br>**phenotype**<br>**in**<br>***C. elegans*** | Mouse:<br>Homozygote<br>has diverse<br>defects;<br>secondary<br>infertility,<br>decreased<br>glucose<br>uptake, mild<br>anemia.<br>Worm:<br>impaired<br>dauer<br>formation. | Ctl2 KO<br>causes<br>progeria,<br>decreased<br>lifespan | Overexpressi<br>on induces<br>resistance to<br>oxidative<br>stress and<br>increased<br>lifespan | Prdx-2 KO<br>Worm:<br>sensitive to<br>oxidation.<br>decreased<br>lifespan. | Trx ko worms:<br>reduced lifespan,<br>sensitive to<br>oxidative stress<br>(paraquat) | Not tested |
| **Homology** | 22.14% ID,<br>(to FOXO3) | 60.53 ID to<br>ctl1 | 55.35 to<br>SOD1 | 72.86 ID to<br>prdx 6 | 33.91% ID to trx1 | 45.10% ID to glrx3 |
| **References** | (90, 91) | (92) | (93) | (94, 95) | (96-98) | (99) |
| **Protein** | *Glutathione S-transferase* | *NAD-dependent protein deacetylase (SIR)* | *Dual specificity mitogen-activated protein kinase kinase* | **Nuclear respiratory factor 1 (NRF1)** | *Dual oxidase 1* | **D-beta-hydroxybutyrate dehydrogenase** |
| **Function** | Antioxidative Enzyme | reduction of the 'Lys-16' acetylation of histone H4 | Kinase activity involved in oxidative stress | mitochondrial DNA transcription and replication | N/A | Oxidative Enzyme |
| **Human** | GSTA 1-5/<br>GSTM 1-5 | SIRT 1-7 | MAP2K1 | NRF1 | Oxidative Enzyme | BDH1 |
| **Murine** | GSTA 1-7/<br>Gstm 1-4 | Sir 1-7 |  | Nfr1 | DUOX1 | Bdh1 |
| ***C. elegans*** | Gst 1-44 | Sir 2.1-2.4 | Mek 1, 2 | Skn 1 | Duox1 | N/A |
| **Location human vs *C. elegans*** | Cytosol vs Cytoplasm and mitochondria | Nucleus, mitochondria and cytoplasm vs Nucleus and cytoplasm | Nucleus and mitochondria vs Cytoplasm | cytoplasm vs Nucleus and mitochondria | bli-3 | Mitochondria |
| **Enzymatic activity** | 121 nmol/min/mg in human<br>N/A for *C. elegans* | N/A | N/A | N/A | Plasma membrane vs plasma membrane | N/A |
| **Disruption phenotype in *C. elegans*** | RNAi-mediated knockdown causes an increase in Mn<sup>2+</sup>-mediated dopaminergic CEP neuron degeneration | Reduces the longevity RNAi-mediated depletion results in an increase of 'Lys-16' acetylation of histone H4 (H4K16ac) | defects in egg laying infertility Reduced lifespan with stress. No obvious phenotype in absence of stress | RNAi-mediated knockdown causes an increase in Mn<sup>2+</sup>-mediated dopaminergic CEP neuron degeneration and a reduction in expression levels of glutathione S-transferase *gst-1* | RNAi-mediated knockdown + proline ROS production, reduces the expression of *skn-1* and reduces longevity | N/A |
| **Homology** | 29.28% ID to GSTA1 | 28.17% ID to SIR1 |  | 12.20% ID to NRF1 | 32.78% | N/A |
| **Refs** | (100, 101) | (102-104) |  | (101, 105, 106) | (107, 108) |  |

| **Protein** | **Xanthine Oxidase** | **NO Synthase** | **Cytochrome P450** | **Hydroxyacid oxidase 1** | **SDH (Succinate Dehydrogenase)** |
| --- | --- | --- | --- | --- | --- |
| **Function** | Oxidative Enzyme | Oxidative Enzyme | Oxidative Enzyme | Oxidative Enzyme | Oxidative Enzyme |
| **Human** | XDH | NOS 1,2,3 | CYP (Multiple) | HAO1 | SDHA, SDHB, SDHC, SDHD |
| **Murine** | Xdh | Nos 1,2,3 | Cyp (multiple) | Hao1 | Sdha, Sdhb, Sdhc, Sdhd |
| ***C. elegans*** | xdh-1 | Unknown | cyp-(23-37) | Unknown | sdha-1, sdhb-1, sdhd-1 |
| **Location human vs *C. elegans*** | Peroxisome and Extracellular region vs Cytosol |  | ER vs Unknown |  | Mitochondria VS Mitochondria |
| **Enzymatic activity** | Rat: 4.4 ccm oxy uptake/unit time/unit weight of intestine | Human: .5-2.5 pmol/min/mg (brain) Mouse: Normalized to 1 | Multiple | --- | 1.45 ferricyanide reduced/min.mg wetwt. (Euthyroid) |
| **Disruption phenotype in *C. elegans*** | Unavailable |  | Unavailable |  | Unavailable |
| **Homology** | 46.54% |  | 28.97% |  | 69.12% |
| **Refs** | (109) |  | (110-113) |  | (114) |

**Table S2:** Amino acid composition of UNC-68 and Human RyR1.

| UNC-68 (*C. elegans*) |  |  | RyR1 (Human) |  |
| --- | --- | --- | --- | --- |
| Number of amino acids | 5187 |  | 5038 |  |
| Molecular weight | 589102.13 |  | 565175.53 |  |
| **Amino acid composition:** | Number | % | Number | % |
| Ala (A) | 333 | 6.4 | 384 | 7.6 |
| Arg (R) | 265 | 5.1 | 294 | 5.8 |
| Asn (N) | 216 | 4.2 | 170 | 3.4 |
| Asp (D) | 302 | 5.8 | 253 | 5 |
| Cys (C) | 90 | 1.7 | 100 | 2 |
| Gln (Q) | 235 | 4.5 | 205 | 4.1 |
| Glu (E) | 417 | 8 | 476 | 9.4 |
| Gly (G) | 305 | 5.9 | 363 | 7.2 |
| His (H) | 133 | 2.6 | 133 | 2.6 |
| Ile (I) | 281 | 5.4 | 203 | 4 |
| Leu (L) | 533 | 10.3 | 558 | 11.1 |
| Lys (K) | 314 | 6.1 | 228 | 4.5 |
| **Met (M)** | **179** | **3.5** | **145** | **2.9** |
| Phe (F) | 251 | 4.8 | 207 | 4.1 |
| Pro (P) | 194 | 3.7 | 271 | 5.4 |
| **Ser (S)** | **376** | **7.2** | **296** | **5.9** |
| Thr (T) | 256 | 4.9 | 231 | 4.6 |
| Trp (W) | 61 | 1.2 | 63 | 1.3 |
| Tyr (Y) | 155 | 3 | 141 | 2.8 |
| Val (V) | 291 | 5.6 | 317 | 6.3 |
| Pyl (O) | 0 | 0 | 0 | 0 |
| Sec (U) | 0 | 0 | 0 | 0 |

## Acknowledgments

This work was supported by grants from the NIH to ARM (T32HL120826, R01HL145473, R01DK118240, R01HL142903, R01HL061503, R01HL140934, R01AR070194).

## Author contributions

HD, FF and ARM designed experiments, analyzed data and edited/wrote the paper. AL, XL, QY, SR, RO, and PS designed experiments and analyzed data.

